# A role for caveolar proteins in regulation of the circadian clock

**DOI:** 10.1101/2022.10.10.511681

**Authors:** Sachini Fonseka, Benjamin D. Weger, Meltem Weger, Nick Martel, Thomas E. Hall, Shayli Varasteh Moradi, Christian H. Gabriel, Achim Kramer, Charles Ferguson, Manuel A. Fernández-Rojo, Kirill Alexandrov, Oliver Rawashdeh, Kerrie-Ann McMahon, Frederic Gachon, Robert G Parton

## Abstract

Caveolae are specialized invaginations of the plasma membrane that are formed by the co-assembly of caveolin integral membrane proteins and a cytoplasmic cavin coat complex. Previous work has proposed an interaction of the cavin coat protein, CAVIN3, with the key circadian clock protein, PER2. Here we show that cavin proteins can play a role in the regulation of the circadian clock by external stimuli. Loss of *Cavin1* in mice caused a shortening of the free-running period of locomotor activity. CAVIN1 and CAVIN3 were found to play a central role in core clock dynamics with either cavin protein directly interacting with PER2 and their perturbation leading to significant disruption in core clock mRNA expression and CRY1 protein oscillation. In cells, association of cavins and PER2 was increased upon caveola disassembly caused by oxidative stress or by calcium influx, stimuli linked to circadian clock regulation. We thus propose that the caveola system can play a modulatory role in circadian regulation through the cavin proteins.

## Introduction

Caveolae are nanoscale sized specialized domains on plasma membranes identified by their unique flask-shape morphology and are particularly enriched in endothelial cells, adipocytes, fibroblasts, and muscle cells whilst rarer in other cell types (Parton, 2018). Caveolae are considered to play an important role in cell signalling, lipid metabolism and trafficking, and provide essential mechanosensing and protective capacity to the membrane (Lamaze et al., 2017, Parton et al., 2020a, Parton et al., 2020b, Sinha et al., 2011, Torrino et al., 2018, Matthaeus et al., 2020, Boscher and Nabi, 2012). The defining components of this membrane domain are the integral membrane caveolin proteins (CAV1-3) and the cavin family of cytosolic coat proteins (CAVIN1-4). Unlike caveolin, cavins are soluble cytosolic proteins that are recruited to the membrane to generate and/or stabilize caveolae. These peripheral membrane proteins have lipid-binding activity and, independent of caveolae, co-associate to form trimers consisting of either two or three CAVIN1 proteins and one other cavin protein (Gambin et al., 2014, Ludwig et al., 2013, Kovtun et al., 2014, Gambin et al., 2013).

The dynamic process of caveola disassembly was first demonstrated under mechanical stress (Sinha et al., 2011). Evidence in recent years suggests that caveola disassembly and cavin coat dissociation can also transmit signals and so act as a membrane sensor (McMahon et al., 2019, Parton et al., 2020b). An example of cavin signalling is the UV-induced release of CAVIN3 triggering apoptosis through direct interaction with the phosphatase PP1α (McMahon et al., 2019). The stimuli-dependent redistribution of cavins from caveolae prompted the proposal of a caveola signal transduction hypothesis, where released cavins could transduce signals from caveolae to effectors in the cytoplasm and nucleus to modulate certain stress responses. Our previous studies involving Amplified Luminescent Proximity Homogenous Assay (AlphaScreen) screening (McMahon et al., 2019) showed an interaction between CAVIN1-3 and the components of the circadian clock PER2 and CRY2. This result supported previous observations suggesting an interaction between the circadian clock caveolae associated signalling (Schneider et al., 2012). However, the cellular context of this interaction and the role caveolae and the cavin proteins play in the molecular clock remained unknown.

The circadian clock is an endogenous timing system consisting of a central pacemaker in the suprachiasmatic nuclei (SCN) of the hypothalamus and of peripheral tissue clocks that orchestrates most aspects of physiology and behaviour. Present in virtually all mammalian cells, rhythmicity in gene expression is generated by a molecular machinery consisting of interlocked transcriptional and translational feedback loops in which the transcription factor BMAL1 (encoded by the *Arntl* gene) plays a critical role. BMAL1 acts on E-box elements on clock-controlled genes (CCGs) including Period (*Per1, Per2, Per3*) and Cryptochrome (*Cry1, Cry2*) genes. PER-CRY complexes can then enter the nucleus and repress the activity of BMAL1. As PER and CRY proteins become degraded through ubiquitin-dependent pathways, BMAL1 activity is restored, and a new cycle can start (Patke et al., 2020). While this process is cell autonomous, the cellular tissue clocks are synchronized by signals originating from the SCN or systemic signals including feeding cues and body temperature (Koronowski and Sassone-Corsi, 2021, Reinke and Asher, 2019). Therefore, we hypothesized that activation of caveola-associated signalling could play a role in the synchronization of the circadian clock by external stimuli.

We now show that both CAVIN1 and CAVIN3 are required for stable core clock gene expression. Strikingly, endogenous CRY1 protein oscillation was significantly disturbed in cells overexpressing either *CAVIN1* or *CAVIN3*. External cues that released cavins from the plasma membrane including oxidative stress and calcium influx, increased the interaction between CAVIN1 or CAVIN3 with PER2. Moreover, PER2 stability was influenced by both CAVIN1 and CAVIN3. Together with characterization of the circadian rhythm of locomotor activity of the *Cavin1* knockout (KO) mouse showing a shorter free-running period length, our results provide evidence that CAVIN1 and CAVIN3 can play a role in regulating the core clock machinery and thus the synchronization of the circadian clock by external stimuli.

## Results

### *Cavin1* KO mice show a shorter circadian free-running period

The role of cavins in the circadian system was first assessed by recording circadian locomotor activity of *Cavin1* KO mice. In this paradigm, single housed male and female mice had unlimited access to a running wheel in light-controlled cabinets. Male and female *Cavin1* KO mice in the C57BL/6 genetic background along with gender- and age-matched wild-type (WT) controls were exposed to a 12:12 hr light:dark (LD) cycle for at least two weeks. Under LD conditions, total daily activity levels and onset of activity were not significantly different between genotypes (Fig 1A). The mice were then allowed to free run in constant darkness (DD) for at least 3 weeks (Fig. 1A). In DD, *Cavin1* KO mice retained rhythmicity. However, the free running period, a fundamental intrinsic characteristic of the central circadian clock, was slightly but significantly shortened in both female (23.54 ± 0.06 h) and male (23.60 ± 0.03 h) KO compared to control (female 23.70 ± 0.02 h and male 23.74 ± 0.03 h) mice (Fig. 1B). This effect on the circadian locomotor rhythm in the *Cavin1* KO mice suggests that CAVIN1 plays a role in the regulation of the circadian period and likely modulates the expression of circadian clock genes.

**Figure 1.**
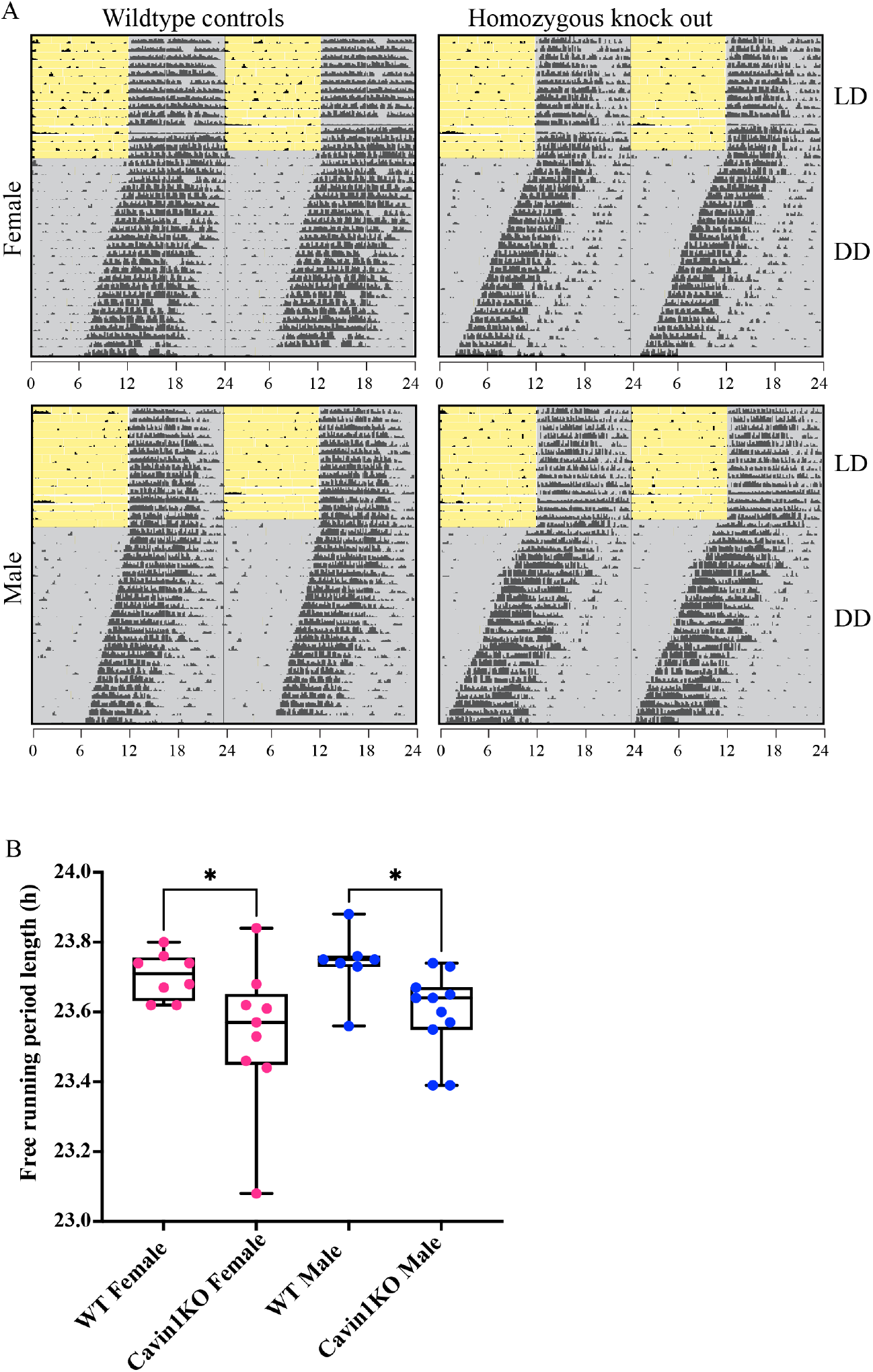
The free-running period length of the Cavin1 KO mouse. **A**| Wheel-running activity was recorded for WT and Cavin1 KO mice under 12h-light-12h-dark cycles (LD) and then in constant darkness (DD). Representative double-plot actograms from males and females of each genotype are shown (8 WT females, 9 KO females, 7 WT males and 11 KO males). **B**|Average free-running period length of mice. * p-value <0.05 Kruskal Wallis test with Dunns multiple comparison tests. Black bars underneath scatter plot presents minimum to maximum box and whisker plots with individual data points shown in colour.

### Role of *Cavin1* and *Cavin3* in core clock gene expression and CRY1 protein oscillation

To test if the reduction in period length observed in the *Cavin1* KO mice is associated with changes in the molecular clockwork, we studied clock gene mRNA expression in genome edited mouse fibroblasts (3T3-L1) lacking *Cavin1* or *Cavin3*. The KO cell lines were validated by quantitative RT-PCR and western blot (Fig. 2A & B). As previously described, the absence of CAVIN1 resulted in a concurrent loss of the CAVIN3 protein, whereas the loss of CAVIN3 had no effect on the expression levels of CAVIN1 (Fig. 2C) (Hill et al., 2008, Hayer et al., 2010). mRNA was collected from unsynchronized cells to observe any overall changes to the core transcription and translational feedback loop (TTFL) and accessory loops. To assess the downstream effect on circadian transcriptional output, we looked at the mRNA expression level of the D-Box Binding PARbZIP transcription factor (*Dbp*), a direct and well described target of BMAL1 (Ripperger and Schibler, 2006, Ripperger et al., 2000). In *Cavin1* KO cells, there was a significant decrease in *Per2, Cry2*, and *Dbp* mRNA while *Cry1* mRNA was increased (Fig. 2D). In slight contrast, in *Cavin3* KO cells, only *Per2* and *Dbp* mRNA levels were significantly decreased. Overall, these findings suggest that loss of *Cavin1* or *Cavin3* leads to the perturbation in the circadian core clock machinery, indicating a role in regulating the function of the molecular clock.

**Figure 2.**
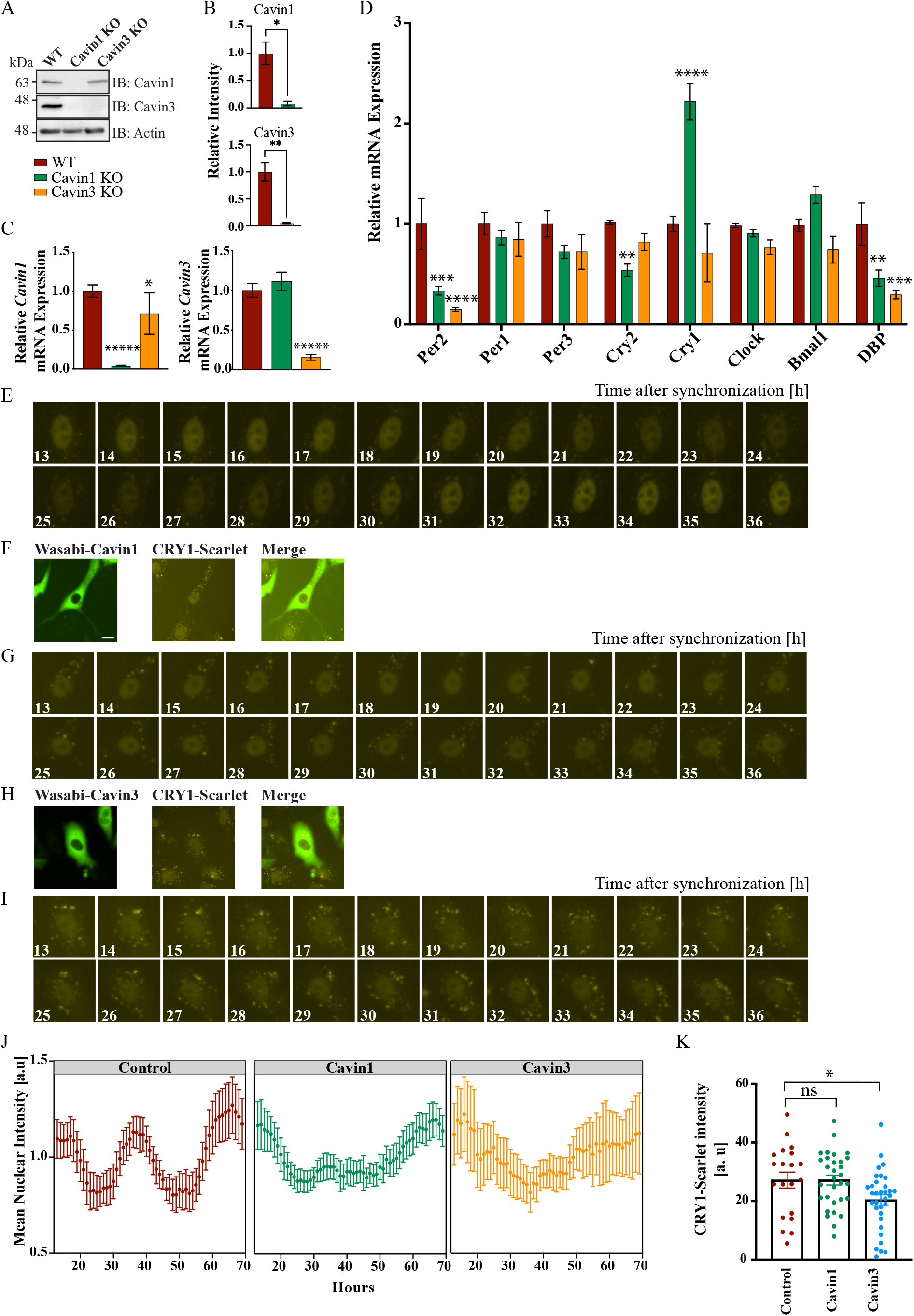
Altered levels of Cavin1 or Cavin3 disturb expression of several core clock genes and disrupts CRY1 oscillation. **A & B**|Validation of Cavin1 and Cavin3 KO 3T3L1 cell lines by immuno-blot of CAVIN1 and CAVIN3. Immunoblots representative of n=3 with ACTIN used as a loading control. Densitometry analysis immunoblots shows KO of protein. Data represents mean± SEM, ** p-value <0.01 and * p-value <0.05 from student’s t-test analysis. **C**| RT-PCR of Cavin1 and Cavin3 in KO 3T3LI cells. Data presented as mean ± SEM. *****p-value <0.00001, and * p-value <0.05 from one-way ANOVA analysis with Turkey’s multiple comparison test. **D**| Per1-3, Cry1-2, Clock, Bmal1 and Dbp mRNA levels in WT, Cavin1 KO, Cavin3 KO 3T3L1 cells as measured by qRT-PCR. RT-PCR data represents n=3 with tubulin used to normalize. Data presented as mean ± SEM. ****p-value <0.0001, *** p-value <0.001, ** p-value <0.01 and * p-value <0.05 from two-way ANOVA analysis comparing to control with Dunnett’s multiple comparison test. **E**|Representative image of nuclear fluorescence in a single CRY1-mScarlet-I knock-in U-2 OS cell 13-36 hrs post-synchronization. **F**| CRY1-mScarlet-I cells transfected with mWasabi-Cavin1 and (**G**) the nuclear monitoring of that selected cell. **H**| CRY1-mScarlet-I cells transfected with mWasabi-Cavin3 and (**I**) the nuclear monitoring of that selected cell. **J**|Background subtracted fluorescent intensities were normalized by dividing intensities by mean intensities of the respective time series. Nuclei were monitored over 3 days post synchronization in cells transfected with either mWasabi-Cavin1 or mWasabi-Cavin3 and non-transfected controls. The first 12 hrs are not shown. Data presented as mean ± SEM, from 20-30 cells measured over two independent experiments. **K**|The total mean nuclear intensity over 3 days in control cells and cells transfected with either mWasabi-Cavin1 or mWasabi-Cavin3. Bar chart represents the mean ± SEM, with individual data overlaid in colour, *p-value <0.05 from a one-way ANOVA compared against the control with Turkey’s multiple comparison test.

To investigate this observation further, we utilized the newly developed osteosarcoma (U-2 OS) *CRY1*-mScarlet-I knock-in cells that allow the live monitoring of CRY1 protein dynamics in single cells (Gabriel et al., 2021). *CRY1*-mScarlet-I cells were transfected with mWasabi-*Cavin1* or mWasabi-*Cavin3*, synchronized with dexamethasone, and imaged every hour for 69 hours (13-36 hrs post-synchronization are shown in the montage Fig. 2E-I; mWasabi reporter control in Supp. Fig. 2B). mScarlet fluorescence was measured in individual nuclei over the entire time course (Supp. Fig. 2A). Fluorescent intensity of nuclear CRY1-mScarlet-I in control cells showed a robust circadian rhythm, whilst CRY1-mScarlet-I in cells overexpressing either Wasabi-CAVIN1 or Wasabi-CAVIN3 displayed a dampened rhythm (Fig. 2J). Furthermore, the overall intensity of CRY1-mScarlet-I was significantly reduced in cells expressing Wasabi-CAVIN3 but not Wasabi-CAVIN1 (Fig. 2K). Collectively, these results suggest that both CAVIN1 and CAVIN3 play a role in CRY1 protein oscillation and CAVIN3 specifically influences protein abundance.

### CAVIN1/CAVIN3 interactions with PER2 are regulated by oxidative stress

Our previous studies involving AlphaScreen analyses (McMahon et al., 2019) and work by Schneider and colleagues (Schneider et al., 2012) corroborate the interaction between PER2 and CAVIN3. However, CAVIN3 localizes predominantly to surface caveolae whereas PER2 localizes to the cytoplasm and nucleus, raising the question of how the two proteins can associate. We therefore hypothesized that an interaction between cavins and PER2 could occur in response to stimuli that release cavins into the cell. As previously shown (McMahon et al., 2019), cells treated with hydrogen peroxide (1 mM H_2_O_2_, 60 min) to induce oxidative stress redistributed CAVIN1 and CAVIN3 into the cytoplasm (Fig. 3A). To evaluate the association between PER2 with CAVIN1 or CAVIN3 in response to oxidative stress, an *in situ* proximity ligation assay (PLA) was performed. This method allows the detection of protein-protein associations with high sensitivity and specificity (see PLA controls Supp. Fig. 3A). Human epithelial cells (HeLa) expressing *PER2*-MYC were treated with H_2_O_2_ and association with endogenous CAVIN1 and CAVIN3 was measured. A GFP reporter was co-transfected to allow detection of MYC transfected cells. The PLA assay was performed with antibodies against MYC and endogenous CAVIN1 or CAVIN3 (Fig. 3B), with GFP only transfected cells used as a negative control (Supp. Fig. 4A). In untreated cells, a low level of association of CAVIN1 and CAVIN3 with PER2 was observed as indicated by a low but significant PLA signal. Upon H_2_O_2_ treatment this association was significantly increased (Fig. 3B and C). No increase in PLA signal was observed in the negative control (Fig. 3C, Supp. Fig. 4A). These results demonstrate that the association between CAVIN1 and CAVIN3 with PER2 is dynamic and modulated by oxidative stress.

**Figure 3.**
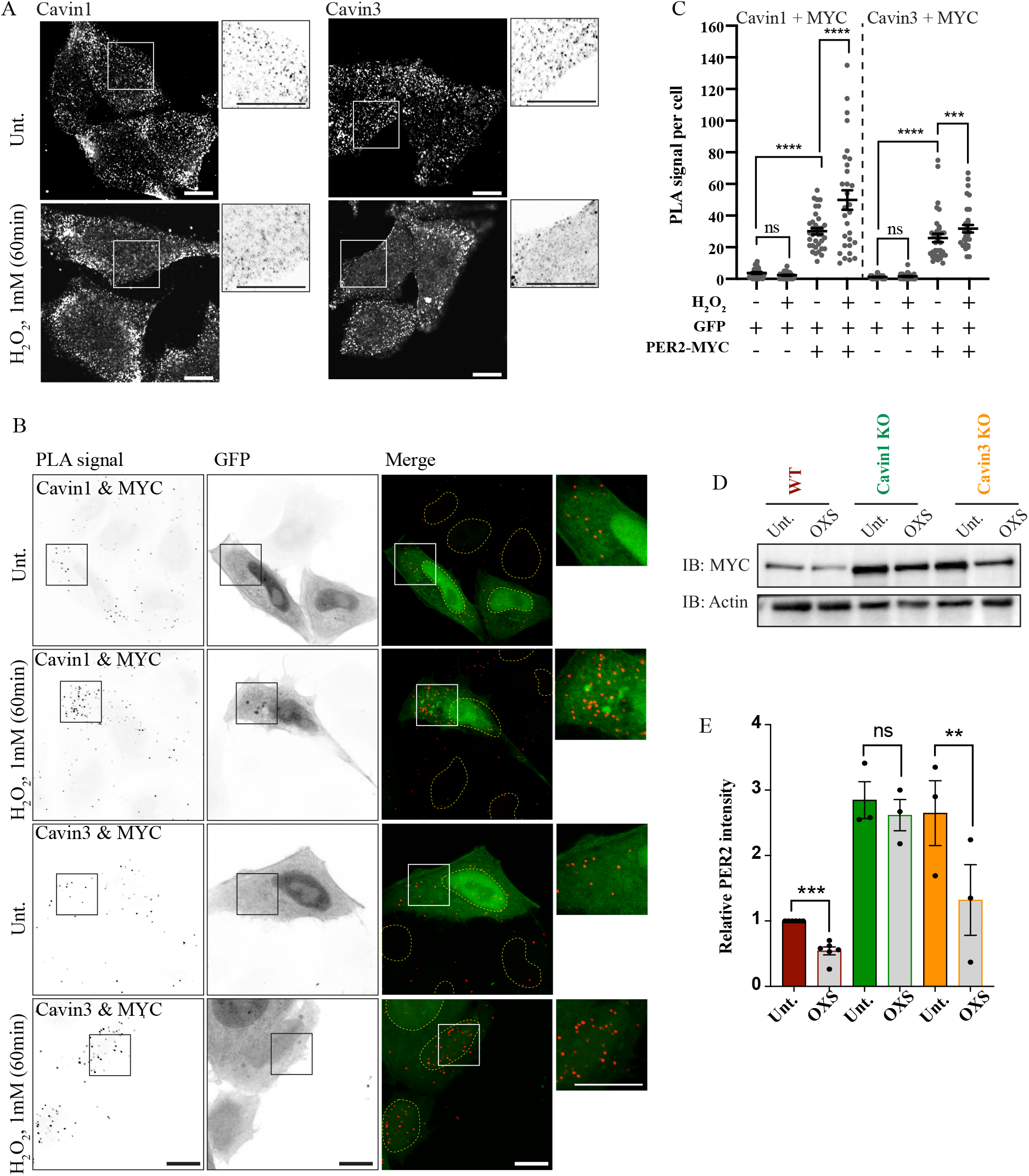
The interaction between CAVIN1 and CAVIN3 with PER2 is regulated by oxidative stress. **A**| Representative immunofluorescence images of endogenous CAVIN1 and CAVIN3 in untreated (Unt.) or oxidative stress conditions (1mM H_2_O_2_, 60min) in HeLa cells. Inverted enlarged images showing the distinct punctate and diffused labelling in boxed areas. **B** |Cells co-transfected with PER2-MYC and GFP reporter, followed by in situ PLA detection of the interaction between PER2-MYC and endogenous CAVIN1 or CAVIN3 in untreated or oxidative stress conditions. Representative images of PLA signals (inverted image, black puncta), GFP signal and merged images (PLA as red puncta). Dashed circles indicate the outline of the nucleus. Enlarged images showing the distinct puncta labelling in boxed areas. **C**|Quantification of PLA signals for the CAVIN1/PER2-MYC and CAVIN3/PER2-MYC interactions upon oxidative stress, a total of 30-40 cells from three independent experiments. Cells transfected with a GFP reporter only was used as a negative control. *** p-value <0.001, ** p-value <0.01 and * p-value <0.05 (one-way ANOVA with Turkey’s multiple comparison test.) **D**| WT and CAVIN1 and CAVIN3 KO cells expressing PER2-MYC were treated with hydrogen peroxide (1mM, 60 min) to induce oxidative stress. Immunoblots are representative of at least three independent experiments. E| Densitometry analysis of PER2 protein, compared to control untreated. ** p-value <0.001 and *** p-value <0.0001 from pairwise Student t-test. Actin was used as a loading control in all experiments. All microscopy and immunoblot images representative of three independent experiments and all data represents mean ± SEM. Scale bar, 10 μm.

Western blot analysis of the cells under oxidative stress indicated that CAVIN1 and CAVIN3 protein levels were unchanged, suggesting that the increased PLA signal was due to a change in the localization of the cavins and not increased protein abundance (Supp. Fig. 4B). However, PER2 was consistently downregulated after one hour of oxidative stress in control WT cells (Fig., 3D & E. Interestingly this was not observed in *CAVIN1* KO cells, with the protein level remaining unchanged after oxidative stress (Fig. 3D & E. In contrast, *CAVIN3* KO cells showed PER2 downregulation as in WT cells. This indicates a specific role for CAVIN1 in regulating PER2 levels in response to oxidative stress.

### CAVIN1 and CAVIN3 play a role in regulating PER2 stability

Given the evidence that CAVIN1 and CAVIN3 associate with PER2, we next explored the potential function of this interaction using our genome-edited cell lines. *PER2*-MYC transfected into *CAVIN1* and C*AVIN3* KO HeLa cells was significantly upregulated compared to WT controls (Fig. 4A). This was unique for PER2-MYC as other transfected proteins were found to be expressed at similar levels between WT and KO cells (Supp. Fig 5A). PER2 was observed at two molecular weights: a larger much more abundant species at ∼180 kDa and a less abundant ∼135 kDa species, taken to represent two stages of post-translationally modified PER2 (Lee et al., 2001). The higher molecular weight PER2 was significantly upregulated in both KO cells whilst the lower molecular weight PER2 was only increased in *CAVIN3* KO cells (Fig. 4B). Quantification of *PER2* mRNA revealed that the mRNA level was not differentially regulated in the KO cells, further indicating that the protein changes observed are due to modified stability (Supp. Fig. 5B). Next, we co-transfected *PER2*-MYC into WT cells alongside GFP-*Cavin1* and GFP-*Cavin3*. Complementing the KO data, PER2 was significantly downregulated when co-expressed with GFP-CAVIN3 (Fig. 4C & D). However, co-expression with GFP-CAVIN1 did not affect the abundance of PER2 (Fig. 4C & D). Together these data suggest that both CAVIN1 and CAVIN3 play a role in regulating PER2 stability with specific nuances for each cavin.

**Figure 4.**
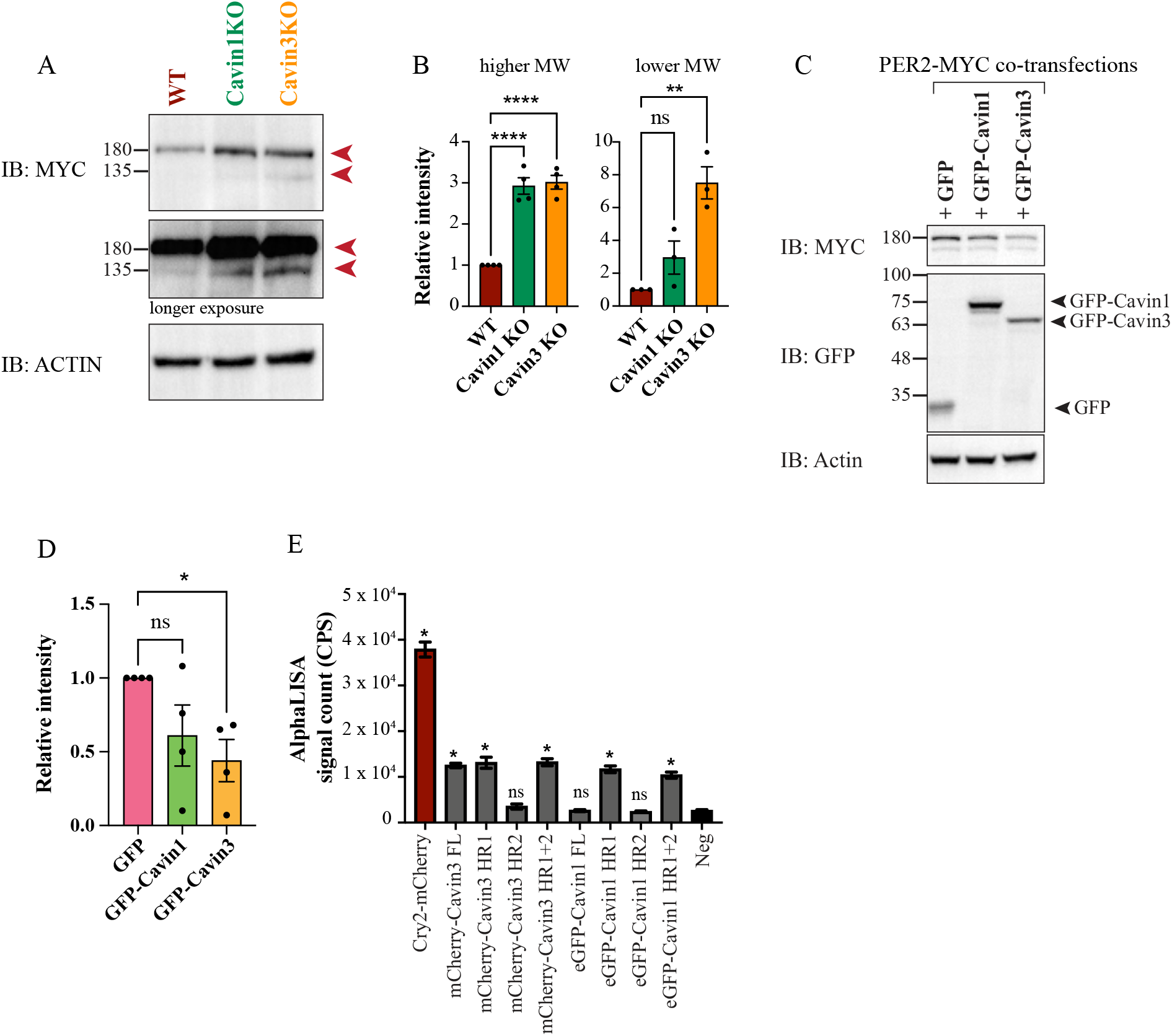
CAVIN1 and CAVIN3 regulate PER2 stability. **A**| WT, CAVIN1 KO and CAVIN3 KO HeLa cells were probed for PER2 24 hours post transfection with PER2-MYC. Immunoblots are representative of three or four independent experiments. A higher and lower molecular weight PER2 species was observed, indicated by red arrows. **B**| Densitometry analysis of high and low MW PER2, compared to control levels in the same experiment. **** p-value <0.0001 and ** p-value <0.01 from one-way ANOVA analysis comparing to control with Turkey’s multiple comparison test. **C**| WT Hela cells co-transfected with PER2-MYC and GFP, or GFP-CAVIN1 or GFP-CAVIN3 and probed for PER2. Immunoblots are representative of four independent experiments. **D**| Densitometry analysis of PER2, compared to control levels. * p-value <0.01 from one-way ANOVA analysis comparing to control with Turkey’s multiple comparison test. **E**| AlphaLISA interaction analysis of CAVIN1 and CAVIN3 full length (FL) and HR domains (HR1, HR2 and HR1+ HR2) with full length PER2. Interaction with CRY2 was a positive control and interaction with FK506 binding protein (FKBP) as a negative control. An arbitrary low threshold of 10,000 CPS of AlphaLISA signal was selected as the cut off for a positive interaction.

To further interrogate the nature of the cavin-PER2 interaction, we assessed the binding affinity of specific CAVIN3 and CAVIN1 domains to PER2 using the *in vitro Leishmania tarentolae* expression system (LTE) and AlphaLISA assay. Human *PER2* and *CRY2* cDNA from the human ORFeome library were cloned into LTE cell free expression vectors. Domain truncated mutant versions of *CAVIN1* (full length (FL), HR1, HR2 and HR1+HR2) and *CAVIN3* (HR1, HR2 and HR1+HR2) were used to establish the specific binding domains (Supp. Fig. 5C). The co-expression efficiency of eGFP or mCherry tagged PER2 with mutant CAVIN1 and CAVIN3 were established in an LTE system (Supp. Fig. 5D). The PER2-CRY2 interaction was used as a positive control and interaction with FK506 binding protein (FKBP) as a negative control. CAVIN3 FL interacted with PER2, as did CAVIN3 HR1 and CAVIN3 HR1 + HR2 but not CAVIN3 HR2 (Fig. 4E). This indicates a specific affinity for the HR1 domain of CAVIN3. Similarly, the HR1 domain of CAVIN1 interacted with PER2 as seen by a positive signal with CAVIN1 HR1 and with CAVIN1 HR1+HR2. However, CAVIN1 FL which contains both HR domains and the disordered regions did not show specific interaction with PER2 (Fig. 4E). This may be due to the HR1 domain of CAVIN1 is being potentially masked in the context of the full-length protein. Additionally, CAVIN1 is known to oligomerize with CAVIN3 and therefore the interactions observed in cells could potentially be mediated by CAVIN3 and not through direct protein-protein interaction. This *in vitro* assay further establishes the aforementioned associations and interactions between CAVIN1 and CAVIN3 with PER2 as a direct protein-protein interaction and also narrows down the HR1 domain to be the direct binding partner. Taken together these results suggest that both cavin proteins interact directly with PER2, most likely through the HR1 domains, and negatively modulate PER2 protein stability.

### Influx of intracellular calcium releases CAVIN1 and CAVIN3 from caveolae and upregulates their interaction with PER2

A recent report showed that *in vivo* and *in vitro* contraction of skeletal muscle influences local *Per2* gene expression through a calcium-dependent pathway (Small et al., 2020). Given the high density of caveolae on muscle membranes, we explored the idea that contraction and/or calcium signals can send signals to the core circadian clock through cavin proteins. We first examined endogenous CAVIN1 and CAVIN3 distribution after treatment with ionomycin, a Ca^2+^ ionophore. HeLa cells were treated with ionomycin in Ca^2+^-free media or in 1mM Ca^2+^ media and compared to an untreated control. Both CAVIN1 and CAVIN3 were retained at the plasma membrane in cells treated with ionomycin in Ca^2+^-free media, but rapidly released into the cytoplasm when treated with ionomycin in the presence of 1 mM Ca^2+^ (Fig. 5A). This was not observed in the presence of a cell-permeable Ca^2+^ chelator, BAPTA (Supp. Fig. 6A). A line scan across the plasma membrane showed the intensity of distribution of CAVIN1 and CAVIN3 away from the membrane after ionomycin treatment in the presence of Ca^2+^ (Fig. 5B). Electron microscopy revealed that morphologically-recognizable surface-connected caveolae were significantly reduced (29 ± 6%, mean±SEM, >20 images, 6 different areas, of two independent experiments) in cells treated with ionomycin in the presence of Ca^2+^ relative to ionomycin alone treatment (Supp. Fig. 6B. To determine if the Ca^2+^-mediated redistribution of CAVIN1 and CAVIN3 results in an increased interaction with PER2, we used the PLA assay to test the association between exogenously expressed PER2 and endogenous CAVIN1 or CAVIN3 (Fig. 5C). Cells treated with ionomycin in the presence of Ca^2+^ recorded a significantly higher number of PLA puncta compared to both controls, indicating more association between CAVIN1 and CAVIN3 with PER2 in response to Ca^2+^ influx (Fig. 5D & E, see GFP controls in Supp Fig. 5C & D). These results suggest that changes in Ca^2+^ levels can link caveolae to the core clock machinery through release of cavin proteins that can interact with PER2 and modulate its stability.

**Figure 5.**
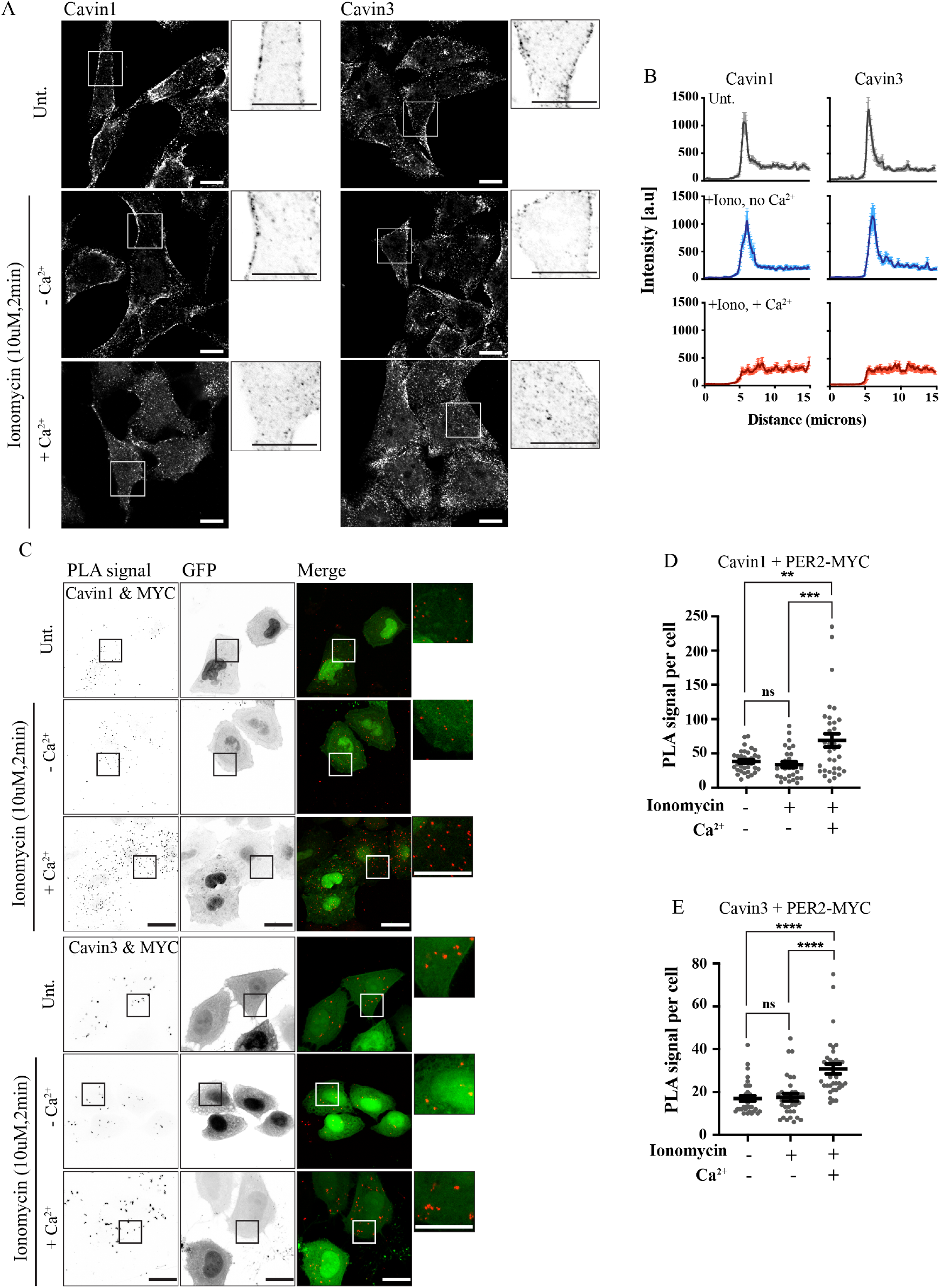
Intracellular calcium releases CAVIN1 and CAVIN3 from the caveolar membranes and upregulates their interaction with PER2. **A**| Representative immunofluorescence images of endogenous CAVIN1 and CAVIN3 in untreated control (Unt.) or cells treated with ionomycin without calcium (10 µM ionomycin no Ca^2+^ media, 2 min) or ionomycin with calcium (10 µM ionomycin in 1mM Ca^2+^, 2 min). Inverted enlarged images showing the distinct punctate and diffused labelling. Scale bar, 10 µm. **B**| A line scan across the cell demonstrating the signal distribution (20 cells quantitated per condition with a 15 µm line scan). **C**| Cells co-transfected with PER2-MYC and GFP reporter, followed by in situ PLA detection of the interaction between PER2-MYC and endogenous CAVIN1 or CAVIN3 in untreated control or ionomycin treatment conditions. Representative images of PLA signals (inverted image, black puncta), GFP signal and merged images (PLA as red puncta). Enlarged images showing the distinct PLA puncta labelling of boxed areas. Scale bar, 20 µm. **D & E**|Quantification of PLA signals for the CAVIN1/PER2-MYC and CAVIN3/PER2-MYC interactions, a total of 30-40 cells from three independent experiments. Cells transfected GFP reporter only were used as a negative control. *** p-value <0.001, ** p-value <0.01 and * p-value <0.05 from one-way ANOVA analysis with Turkey’s multiple comparison test. All microscopy images are representative of three independent experiments and all data represents mean ± SEM).

## Discussion

In this study we used a variety of *in vivo* and *in vitro* models to investigate the role of CAVIN1 and CAVIN3 in the molecular clock. We showed that *Per2* and *Dbp* mRNA were significantly downregulated in both *Cavin1* and *Cavin3* deficient cells, whilst other components were differentially perturbed, suggesting a potential separate function of the cavins. Specifically, the downregulation of *Dbp* mRNA in both our KO cells may indicate perturbed clock function due its transcription being directly controlled by BMAL1/CLOCK (Ripperger and Schibler, 2006, Ripperger et al., 2000). Moreover, we observed that overexpression of CAVIN1 or CAVIN3 severely perturbed the rhythm of CRY1 protein oscillation in the nucleus. The expression pattern of CRY1 is a pivotal component in the autoregulatory loop. CRY1 is thought of as the dominant repressor in the TTFL as it directly binds to CLOCK/BMAL1 to inhibit transcription and is necessary to mediate PER2’s secondary inhibitory effect (Cao et al., 2021, Ye et al., 2014). Consequently, deletion of *Cry1* and *Cry2* genes in mouse leads to an arrhythmic behaviour (Vitaterna et al., 1999, van der Horst et al., 1999). Considering recent single particle evidence that PER2 is almost always in complex with CRY1/2, we can confidently speculate that this perturbation would affect PER2 dynamics (Aryal et al., 2017). The essential role of CRY1 in the molecular clock strongly suggests that overexpression of CAVIN1 or CAVIN3 affects the timing of the repressive loop and ultimately disrupts the core oscillator. Additionally, the stability of PER2 protein was influenced by the level of CAVIN1 and CAVIN3. Exogenous PER2 is consistently elevated in the cells deficient in either *Cavin1* or *Cavin3* and, conversely, co-expression with CAVIN3 led to downregulation of PER2. Potentially the role of PP1α in this interaction is a missing element. We have previously demonstrated that both CAVIN1 and CAVIN3 interact with PP1α directly via the HR2 domain and that CAVIN3 has an inhibitory effect on its activity (McMahon et al., 2019). Modulation of PP1 activity by specific inhibitors or genetic manipulation showed its important role in the regulation of PER2 phosphorylation and degradation, as well as its impact on the regulation of the circadian period (Gallego et al., 2006, Lee et al., 2011, Schmutz et al., 2011). The loss of *Cavin1* and *Cavin3* would relieve the inhibition of PP1αand thus enable its dephosphorylation of PER2 and increased stability, as observed in our results.

At steady state, cavin proteins are predominantly associated with plasma membrane caveolae. Cavin protein levels in the cytosol are maintained at a low level due to ubiquitination and proteasomal degradation (Tillu et al., 2015). We therefore explored the hypothesis that stimuli that redistribute cavins could regulate their interaction with PER2. Recent work from our group demonstrated that UV and oxidative stress release CAVIN1 and CAVIN3 into the cytoplasm (McMahon et al., 2019, McMahon et al., 2021). Here we show that the association between CAVIN1 and CAVIN3 with PER2 is upregulated after oxidative stress. Oxidative stress is known to impact the molecular clock, with high doses of H_2_O_2_ acting as strong synchronisation signal in cultured cells (Tahara et al., 2016) and the sensitivity to reactive oxygen species to be dependent on the cells circadian time due to the rhythmic expression of major antioxidative enzymes (Wilking et al., 2013, Hardeland et al., 2003, Magnone et al., 2014). However, the precise mechanisms by which oxidative stress impacts the clock are unknown. Here we propose a model in which the cavins are primed to relay stress stimuli to the cellular molecular clock through direct interaction with PER2. Interestingly, H_2_O_2_ exposure caused a degradation of exogenous PER2 in control cells and cells lacking *CAVIN3* but had no effect in cells without *CAVIN1* (Figure 3D). Overexpression of both cavin proteins lead to a similar decrease in PER2. This indicates the potential role of CAVIN1 in targeting PER2 stability in response to oxidative stress and its potential role in the synchronization of the circadian clock.

In view of our hypothesis that cavin release from caveolae membranes regulates their interaction with PER2, we wanted to further explore mechanisms that could stimulate caveola disassembly. A recent publication that demonstrated skeletal muscle PER2 expression responds to an intracellular Ca^2+^ influx prompted us to question whether caveolae are involved in this process (Small et al., 2020). Strikingly, a short bout of ionomycin treatment in the presence of Ca^2+^ released cavins from the membrane. Concurrently this led to the significant decrease in caveola density at the membrane (Figure 5C). Calcium reduces phosphatidylinositol 4,5-bisphosphate [PI(4,5)P2] via phosphoinositide phospholipase C (PLC) activation (Allen et al., 1997) and cavins have a medium affinity binding to PI(4,5)P_2_ (Kovtun et al., 2014). Subsequently, mutation in the cavin binding domain with PI(4,5)P_2_ reduces cavin retention at the membrane (Kovtun et al., 2014). Fujita *et al*., demonstrated the cleavage of the local caveolae PI(4,5)P_2_ cluster in response to calcium influx occurs at a faster rate compared to the undifferentiated flat membrane (Fujita et al., 2009). This is potentially due to calcium influx occurring preferentially near caveolae (Isshiki and Anderson, 2003). In addition to increased hydrolyses of PI(4,5)P_2_, Ca^2+^ influx induces a transmembrane movement of PI(4,5)P2 and phosphatidylserine from the inner to the outer plasma membrane (Sulpice et al., 1994). The combination of depleting both PI(4,5)P_2_ and phosphatidylserine from the inner leaflet would be a strong signal for cavin release. Taken together it is clear that these specialized microdomains harbouring cavin proteins are poised Ca^2+^ signal transductors. Similar to oxidative stress, this increase in the cytosolic cavin pool enhanced the association with PER2. The role of intracellular Ca^2+^ in regulating the molecular clock has largely been investigated in the scope of coupling cells in the neuronal pacemaker circuit (Harrisingh et al., 2007; Ikeda et al., 2003; Lundkvist et al., 2005; Noguchi et al., 2017). Our current understanding of the mechanism by which Ca^2+^ signals the TTFL is through calcium dependent CREB phosphorylation and binding to CRE elements to change clock gene expression (Impey et al., 2004; O’Neill et al., 2008; Pulivarthy et al., 2007; Tischkau et al., 2003). Our discovery that Ca^2+^ regulates the CAVIN1-3/PER2 interaction presents a new and unique CREB independent pathway in which Ca^2+^ acts on the molecular clock. The function of this enhanced association is yet to be elucidated. Previous studies by Schneider *et al*., (2012) using *Caveolin1* null mice that lack caveolae but expressed cavins did not find an effect on period length of locomotor activity. Thus, the loss of caveolae in the *Cavin1* null mice may not be a contributing factor to the circadian phenotype observed here, and instead highlights the specific involvement of cavin proteins. The circadian period of activity is a direct readout of the function of the circadian clock of the central circadian pacemaker in the SCN (Hastings et al., 2018). Interestingly, while neurons, the main cellular type in the SCN, are known to be caveolae-deficient, caveolae are abundant in astrocytes (Cameron et al., 1997). Recent evidence supports an important role of the astrocytes in the SCN. The circadian clock in astrocytes is required for normal circadian rhythms (Barca-Mayo et al., 2017, Brancaccio et al., 2017, Tso et al., 2017) and astrocytes are important to generate the rhythm and keep the synchrony of surrounding neurons, particularly through Ca^2+^ signalling (Brancaccio et al., 2017, Patton et al., 2022, Brancaccio et al., 2019). The response of astrocytes to external stimuli involves the intricate relationship between caveolae and fatty acid-binding protein 7 (FABP7) through the formation and function of lipid microdomains (‘rafts’) (Kagawa et al., 2015). Interestingly, FABP7 is rhythmically expressed in astrocytes through direct control by the circadian clock (Schnell et al., 2014, Gerstner et al., 2012), this regulation playing likely an important role in sleep homeostasis (Gerstner et al., 2017, Mang et al., 2016). Therefore, it is tempting to speculate that caveolae could play a role in the synchronization of the circadian clock in response to stresses such as sleep deprivation that is known to induce oxidative stress (Mathangi et al., 2012, Pandey and Kar, 2018, Vaccaro et al., 2020). Future experiments design to define the potential astrocyte-specific role of caveolae in sleep and circadian rhythms would answer this question.

Collectively, our studies demonstrate that the caveola coat proteins CAVIN1 and CAVIN3 are modulators of the molecular clock machinery. CAVIN1 and CAVIN3 directly interacts with the key negative limb component PER2 and, most uniquely, this interaction is regulated by signals that release CAVIN1 and CAVIN3 from caveolae into the cytosol. Both oxidative stress and calcium influx upregulates the association in a dynamic fashion, and thus presents a unique signalling pathway to the molecular clock. Thus, these findings suggest that circadian rhythm regulation is an exciting new component of caveola function and cavin signalling.

## Material and Methods

### Cell culture and reagents

3T3-L1 (CL-173, *Mus Musculus*, fibroblast), HeLa (CRM-CCL-2, *Homo Sapiens*, epithelial) and U-2 OS (HTB-96, *Homo Sapiens*, osteosarcoma) cells were cultured in Dulbecco’s modified Eagle’s medium (DMEM) supplemented with 10% (v/v) fetal bovine serum (FBS), 100 units/ml penicillin and 100 μg/ml streptomycin at 37°C in a humidified atmosphere containing 5% CO_2_. DMEM and trypsin-EDTA solution were purchased from Sigma-Aldrich (Australia). Penicillin/streptomycin was from Invitrogen (USA). FBS was from Life Technologies (Australia). Cells were routinely screened for mycoplasma using a MycoAlert Mycoplasma detection kit (#LT07-418; Lonza). Protease Inhibitor Cocktail Set III. EDTA-free (#539134) was from Merck Millipore (Germany) and PhoSTOP™ Phosphatase Inhibitor Cocktail (#04906837001) was from Roche Diagnostics Australia (Australia). Hydrogen peroxide (H_2_O_2_) solution (30%, w/w in H_2_O) was purchased from Sigma-Aldrich (Australia). Cells were transfected with plasmid DNA using Lipofectamine 3000 (Invitrogen) according to manufacturer’s instruction (∼2 µg DNA per 1 × 10^6^ cells).

### CRISPR-Cas9-based generation of CAVIN1 and CAVIN3 KO 3T3 L1 cells

3T3-L1 *Cavin1* and *Cavin3* KO cells were generated at the Queensland Facility for Advanced Genome editing and Genome Innovation Hub, The University of Queensland. For *Cavin1* KO 3T3L1 cells, three highly specific guide RNAs (gRNAs) were designed using the online program CRISPOR (Concordet and Haeussler, 2018). The gRNA sequences for *Cavin1* were: *gRNA1* ATCAAGTCGGACCAGGTGAA, *gRNA2* GCTCACCGTATTGCTCGTGG and *gRNA3* GTCAACGTGAAGACCGTGCG. For *Cavin3* KO 3T3 L1 cells, four highly specific gRNAs were used. The gRNA sequences for *Cavin3* were: *gRNA1* CTTACTCGAGATCCCGGG, *gRNA2* CAGGCCGTGACTCAGCCGGG, *gRNA3* AGGGCGAGCGACAATGCGCA and *gRNA4* ATATTTCTCCGTGCCAATGG. For CRISPR delivery, synthetic gRNA (CRISPR RNA/trans acting CRISPR RNA duplex, 10 pmol each, IDT) were mixed with spCas9 protein (IDT) in a 1:1 ratio and were transfected into 100,000 3T3L1 cells by electroporation (#MPK1025 Neon transfection kit, Invitrogen, three pulses at 1650 V, 10 ms). Editing efficiency of bulk cell pools was initially confirmed by T7E1 (T7 Endonuclease 1, #M0302S, NEB) assay, followed by single-cell cloning using limited dilution in a 96 well plate. Genomic sequence of isolated clones at the target locus was further confirmed by PCR cloning (E#1202S, NEB) and Sanger sequencing (Australian Genome Research Facility, The University of Queensland). Clones were further validated by Western analysis for protein expression.

### Antibodies

The following antibodies were used: rabbit anti-CAV1 (dilution WB 1:3000) (610060 BD Biosciences), rabbit anti-CAVIN1 (dilution IF/PLA 1:200, 18892-1-AP ProteinTech Group), rabbit anti-CAVIN3 (dilution WB 1:1000, IF/PLA 1:200, 16250-1-AP ProteinTech Group), mouse anti-MYC (dilution WB 1:1000, IF/PLA 1:200, 9B11 Cell Signalling Technology) and mouse anti-ACTIN (1:5000, MAB1501 Merck Millipore). Secondary antibodies were anti-rabbit and anti-mouse HRP (G21040 and G21234 Life Technologies Australia) and goat anti-mouse and anti-rabbit Alexa Fluor (A21235 and A27040 ThermoFisher Scientific).

### Quantitative RT-PCR

Total RNA was isolated using the RNeasy mini kit (QIAGEN) according to the manufacturer’s instructions. 1 µg of total RNA was reverse transcribed with random hexamers using the SuperScript III First-Strand Synthesis System for RT-PCR (Life Technologies). Quantitative PCR was performed in duplicate for three independent cDNA preparations using the SYBR Green Master Mix (Applied Biosystems) on a real-time PCR system (ViiA7; Applied Biosystems). Gene expression data were analysed using the ΔΔCt method and normalized to levels of *Tubulin*. Primers are listed in Table 1.

**Table 1.**
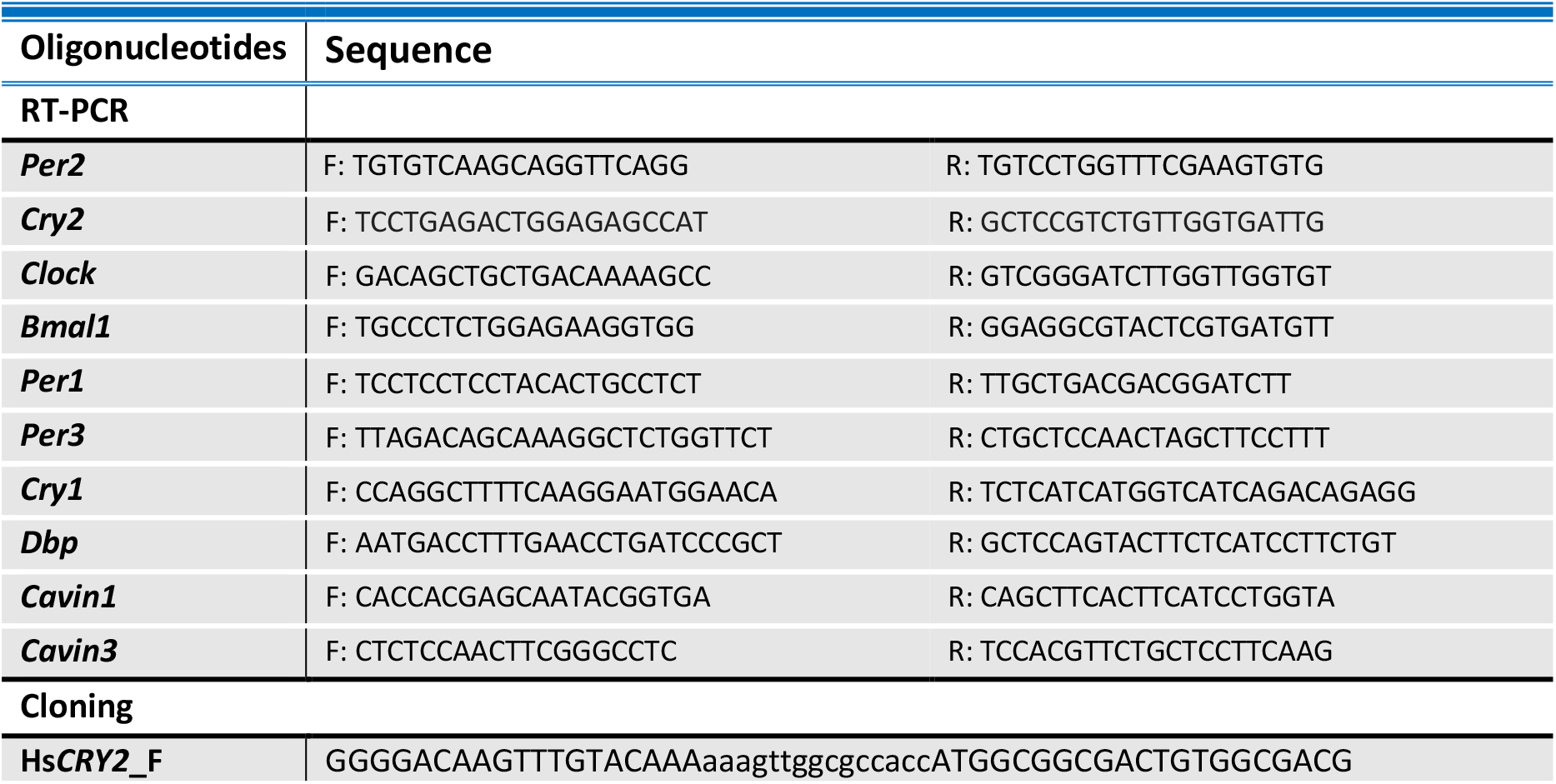
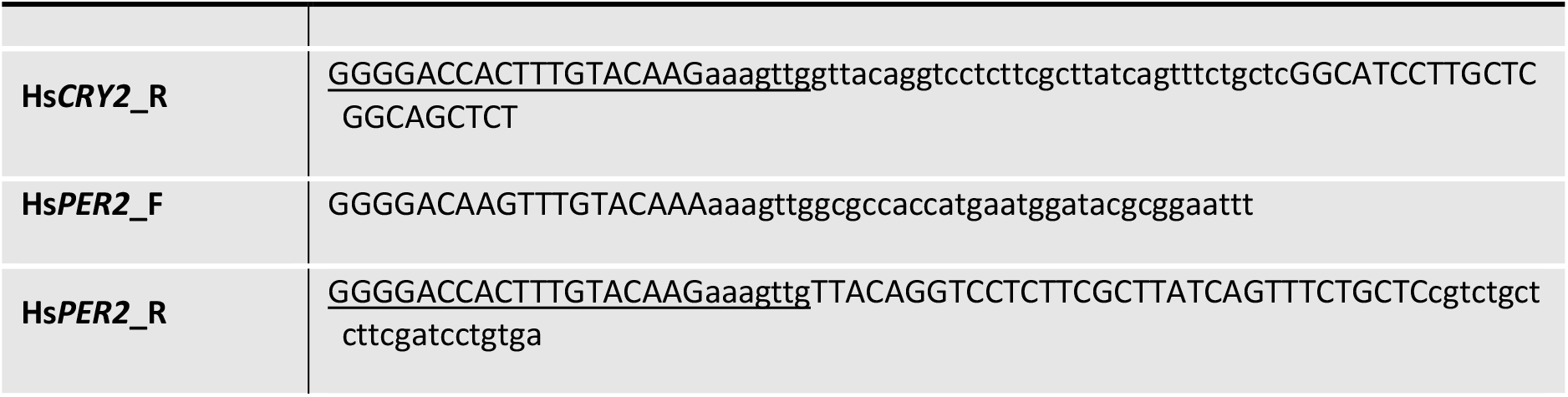
Oligonucleotide sequences.

### SDS PAGE and western blot analysis

Cells were lysed in RIPA buffer (Tris-HCL 50 mM, NaCl 150 mM, SDS 0.1%, Triton X-100 1%, EDTA 5 mM) supplemented with protease (Merck) and phosphatase inhibitor (Roche Diagnostics) cocktail at 4 °C. Protein concentration was determined using a BCA Protein Assay Kit (Thermo Scientific). An equal 20-40 ug of proteins were separated by SDS PAGE and transferred to PVDF membranes (Millipore). Western blots were performed as standard procedures. Detection and quantification of target proteins were carried on the BIO-RAD, ChemiDoc system using horseradish peroxidase (HRP)-conjugated secondary antibodies and ECL detection reagent (Thermo Scientific).

### Cloning

Human *PER2* and *CRY2* cDNA clones from the Human ORFeome were cloned into cell free expression constructs expressing N-terminal eGFP (#5797) or C-terminal mCherrry (#5800) using Gateway Cloning technology. Full sequence identity was confirmed via sanger sequencing using generic M13 forward and reverse, eGFP and mCherry specific primers. The same human cDNA clones were used to develop c-terminal MYC tagged proteins. Briefly, the coding sequence was amplified out with PrimeStar Taq using primers designed with a MYC and kozak sequence (see Table 1). Through Gateway cloning these were synthesised into CMV driven mammalian expression vector. Full sequence identity was confirmed via sanger sequencing using internal primers designed for *CRY2* and *PER2* (see Table 1).

### Immunofluorescence

HeLa cells were plated on glass coverslips at 70% confluence and then fixed in 4% (v/v) paraformaldehyde in PBS for 15min at room temperature (RT). Coverslips were then washed three times in PBS, permeabilized in 0.1% (v/v) Triton x-100 for 5 min, and blocked in 1% (v/v) BSA (Sigma Aldrich) for 30 min at RT. The primary antibodies were diluted in 1% (v/v) BSA solution in PBS and incubated for 1h at RT. Secondary antibodies were diluted 1:500 in 1% BSA and incubated for 1h at RT. After three washes in PBS coverslips were mounted in Mowiol (Mowiol 488, Heochst AG) in 0.2M Tris-HCL, pH 8.5. Images were obtained on a laser-scanning microscope (LSM 510METa, Carl Zeiss, Inc) using 63X oil objective lens. ImageJ software was used for image processing.

### Live nuclear monitoring and analysis

The CRISPR-generated *CRY1*-mScarlet-I knock in cells have been previously described (Gabriel et al., 2020). U-2 OS *CRY*1-mScarlet-I knock in cells were seeding on glass bottom #1.5H μ-slides (80806, Ibidi GmbH, Germany) 8 well plates. Cells were transfected with mWasabi constructs and 24 hours post transfection synchronized with 1µM dexamethasone for 1 hour followed by a warm PBS wash. Imaging was performed on a Nikon Widefield Ti2 equipped with a sCMOS, PCO.edge camera and a live-cell incubator. Image acquisition was done in phenol red free Leibovitz L-15 medium (21083027, ThermoFisher Scientific) supplemented with 1 % FBS, 1:100 PenStrep at 37 °C and 5 % CO2. The following light sources (LEDs) and emission filters were used for the different channels: CFP (cerulean): GFP (mWasabi): excitation 475/28 nm, emission 520/26 nm and RFP (mScarlet-I) excitation 555/28 nm, emission 642/80 nm. Objectives: 40x ApoFluor, NA 0.95, WD 250 μm; 20x Plan Apo, NA 0.8, WD 1 mm. Typically, illumination time for RFP was 300 ms and 20 s for GFP channels. Imaging started 2 h after synchronization with a regular imaging interval of 1 h. Fluorescence data were extracted and z-projected to maximum intensity using ImageJ. TrackMate software was used track mScarlet-I positive nuclei over time and extract mean fluorescence intensities (Tinevez et al., 2017). Individual background fluorescence for every cell at every time point was determined by quantifying the same area of the imaging field from a cell-free image frame of the same experiment. Mean background signal was subtracted. To eliminate cell division outliers, fluorescence values at cell division events were excluded.

### LTE protein expression and AlphaLISA interaction analysis

LTE cell-free protein expression in combination with AlphaLISA^®^ approach (Perkin Elmer, AUS) was performed for pairwise protein interaction analysis of PER2 and CRY2 proteins with CAVIN3 and CAVIN1 protein variants. The coding sequences of human *CAVIN1* (full length (FL), HR1, HR2 and HR1+HR2) and *CAVIN3* (HR1, HR2 and HR1+HR2), *PER2* and *CRY2* were synthesised as Geneblocks (IDT) and inserted into pDONR p2R-p3 (Invitrogen). These entry clones were then recombined into cell-free EGFP or mCherry tagged destination vectors using Multisite Gateway cloning as previously described (Hall et al. 2020). To co-express the protein pairs, eGFP and mCherry tagged plasmids (at the concentration of 30 and 40 ng/μL respectively) were added into 7 μL of supplemented LTE lysate. The reaction was topped up with Ultra-Pure™ water (Invitrogen™, AUS) to the final volume of 10 μL and incubated at 25 ^0^C for 5h. LTE lysate production and protein expression set up is described in detail elsewhere (Varasteh Moradi et al., 2021, Johnston et al., 2019).The quality of co-expressed proteins were evaluated through resolving the protein bands on SDS-PAGE gel (Bolt 4-12%, Invitrogen™) Following the completion of protein expression, the mixture was diluted 25× with buffer A (25 mM HEPES, 50 mM NaCl, 0.1% BSA and 0.01% Nonidet P-40). The AlphaLISA assay was carried out in an Optiplate-384 Plus plate, using anti-GFP AlphaLISA Acceptor and Streptavidin Donor beads. The biotinylated mCherry nanobody (100 nM stock) was added into microplate wells at the final concentration of 4 nM followed by addition of 15 µL of diluted protein mixture (PER2 and targets). The acceptor and donor beads stocks (5 mg/mL) were diluted to 100 µg/mL (5×) in buffer A before adding to the wells. The acceptor beads were then added to each well (5 µL) and the mixture was incubated for 30 min at RT. At last, 5 µL of donor beads were added to the samples under subdued light, mixed gently and incubated for 30 min at RT. All samples were prepared in triplicate. The AlphaLISA signal was detected with Tecan Spark multimode microplate reader using the following setting: Mode: AlphaLISA, Excitation time: 130 ms, Integration time: 300 ms.

### Oxidative stress and ionomycin treatment

For oxidative stress experiments HeLa cells were plated on coverslips to reach 70% confluency overnight and fresh media added one hour prior to treatment. Cells were either left untreated or were exposed to 1 mM H_2_O_2_ (H1009 Sigma Aldrich) in DMEM for 1 hour. For calcium influx experiments cells were treated with one of the following mediums for 2 minutes: PBS (control) or 10 mM ionomycin in PBS (no calcium control) or 10 mM ionomycin in 1 mM Calcium and 1 mM Magnesium chloride in PBS (ionomycin with calcium medium) (AB120116 ABCAM). Immediately after cells were either fixed or lysates prepared as described above. Fluorescent images of ionomycin treated cells were quantified using a line scan. Briefly, a 15 µm line scan across the membrane with the cell boundary placed at 5 µm was used to measure the signal distribution in 30 cells over three independent experiments.

### *In situ* PLA

Detection of an interaction between proteins was assessed using the Duolink II Detection Kit (Sigma-Aldrich) according to the manufacturer specifications. The signal was visualized as a distinct fluorescent spot and was captured on an Olympus BX-51 upright Fluorescence Microscope using a ×60/1.35 oil lens. The number of PLA signals in a cell was quantified in Image J using a Maximum Entropy Threshold and Particle Analysis where 10 cells in each group were analyzed for at least three independent experiments. RGB images were converted to black/white images with the Invert LUT from Image J.

### Electron microscopy

For electron microscopy, HeLa cells were treated with ionomycin as described above and then fixed for epon embedding. Sections were cut perpendicular to the culture substratum. Processing and quantitation of the density of caveolae was performed on over 30 images for each condition captured at a primary magnification of 25kx on a Jeol 1011 transmission electron microscope at 80kV. Six sets of images from two independent experiments were analysed for the presence of caveolae on the apical surface of the cell.

### Animal housing and circadian phenotyping

*Cavin1* KO mice, generated by replacing part of Exon 1 of the *Cavin1* gene with a LacZ/Neo fusion cassette, were obtained from Boston University School of Medicine (Liu and Pilch, 2008). *Cavin1* KO mice and WT controls were maintained according to institutional guidelines (University of Queensland). All animal experiments were approved by the University of Queensland Animal Ethics Committee (IMB/076/18/BREED and SBMS/PACE/416/18). A comparison of free running period length between age matched sibling WT controls and purchased WT controls was done to check that purchased WT controls were a suitable control for these mice circadian behaviour (Supp Fig. 1A). Purchased WT mice were used in experiments due to the breeding difficulty in obtaining KO mice and their WT siblings. Circadian phenotyping of the free-running behaviour was conducted as previously described (Jouffe et al., 2016). Briefly, mice between 8 to 15 weeks of age were single-housed in cages, equipped with running wheels, in ventilated, light-tight cabinets. Inside the cabinet, light (250 lux)/dark cycles were controlled and wheel rotations per unit of time measured by the ClockLab (Actimetrics) computer software. The period was determined using Chi-square periodogram.

### Statistical analysis

All results are presented as mean ± SEM (unless otherwise stated). A combination of Microsoft Excel, ClockLab (Actimetrics), R packages and GraphPad Prism was utilized for data analysis. Western blot densitometry quantification was done on ImageJ software. Student’s t-test and ANOVA was used to compare between two or more groups, respectively with the post-hoc test used noted in each figure legend. P-values < 0.05 were considered significant and indicated by an asterisk (*p<0.05, **p<0.01, ***p<0.001, ****p<0.0001).

## Acknowledgements

This work was supported by the National Health and Medical Research Council of Australia (grants APP1140064 to T.E.H. and R.G.P., APP1150083 and fellowship APP1156489 to R.G.P.; grants APP1125390, APP1140906 to M.T.R. and D.A.S.; fellowship APP1140851 to D.A.S.). The authors acknowledge the use of the Microscopy Australia Research Facility at the Centre for Microscopy and Microanalysis at The University of Queensland. Confocal microscopy was performed at the Australian Cancer Research Foundation (ACRF)/Institute for Molecular Bioscience (IMB) Dynamic Imaging Facility for Cancer Biology with funding from the ACRF.

## Author Contributions

Conceptualization, S.F., K.A.M and R.G.P.; Methodology, S.F., C.F., N.M., T.E.H., M.A.F, B.W., M.W., C.G.; Formal Analysis, S.F., R.G.P, S.V.M.; Writing – Original Draft, S.F., K.A.M. and R.G.P.; Writing – Review & Editing, S.F., K.A.M., T.E.H., O.R., MW, B.D.W, F.G. and R.G.P.; Supervision, K.A., O.R., F.G., A.K. and R.G.P.; Funding Acquisition, T.E.H. and R.G.P.

