## Supplementary Figures for "A role for caveolar proteins in regulation of the circadian clock"

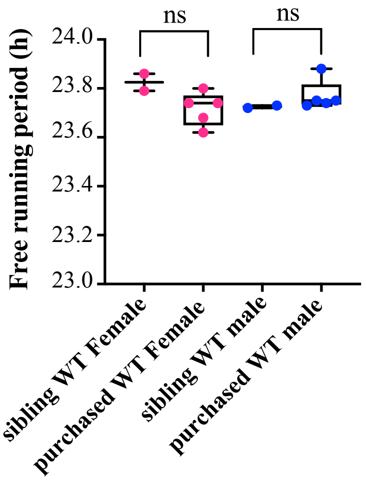


***Supplementary Figure 1*** / ***Free running period length of littermate WT and purchased mice.*** *Wheel-running activity was recorded for age and gender matched sibling WT and purchased WT mice under 12h-light-12h-dark cycles (LD) and then in constant darkness (DD). Average free-running period length of mice in DD is shown; ns, not significant (Kruskal Wallis test with Dunns multiple comparison tests). Black bars underneath scatter plot presents minimum to maximum box and whisker plots with individual data points shown in colour.*

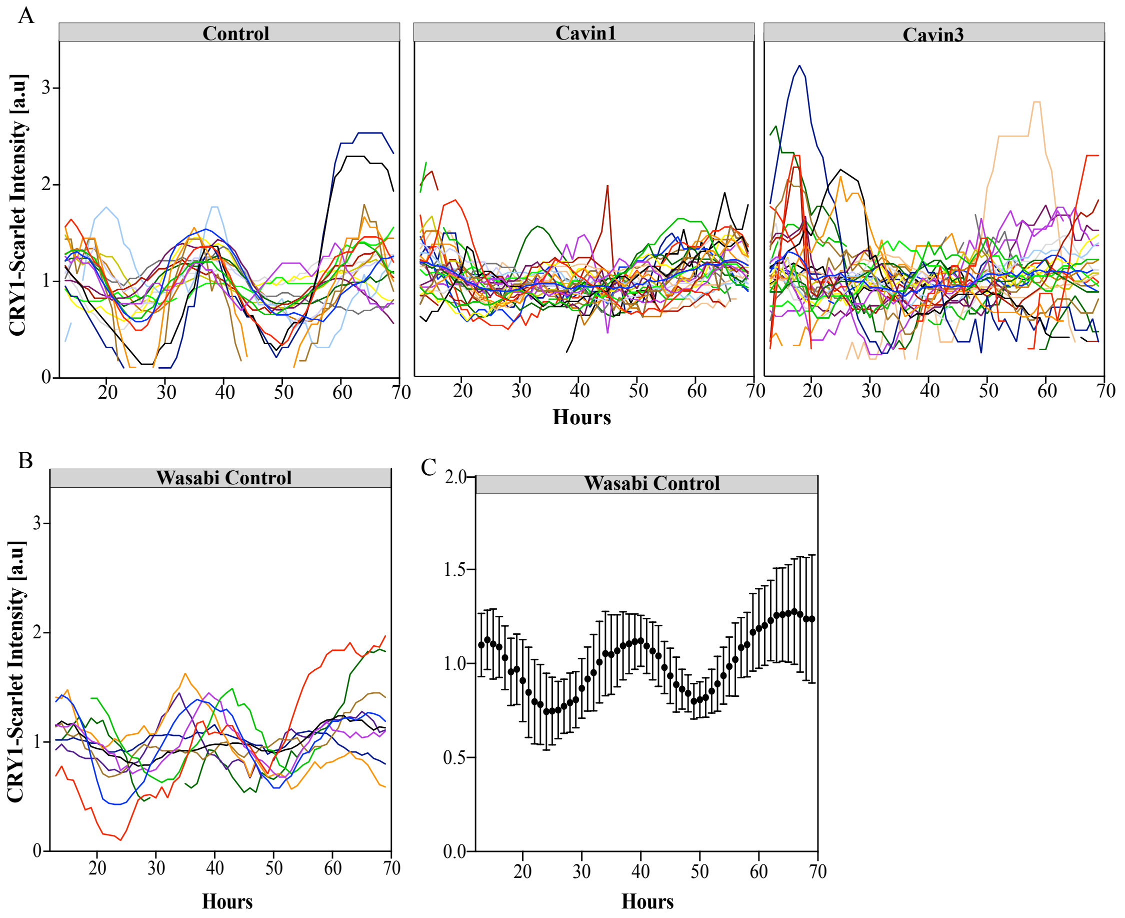


***Supplementary Figure 2*** / ***Individual nuclei traces of CRY1-mScarlet-1 cells. A*** / *Background subtracted fluorescent intensities were normalised by dividing intensities by mean intensities of the respective time series. Nuclei were monitored over 3 days post synchronization in cells transfected with either mWasabi-Cavin1 or mWasabi-Cavin3 and non-transfected controls. The first 12 hours are not shown, 20-30 cells were recorded per condition.* ***B*** / *Individual nuclei traces of CRY1-mScarlet-1 cells expressing mWasabi protein, a transfection/expression control. The first 12 hours are not shown, 10 cells were recorded.* ***C*** /*The mean and standard error of traces shown in Fig 2B.*

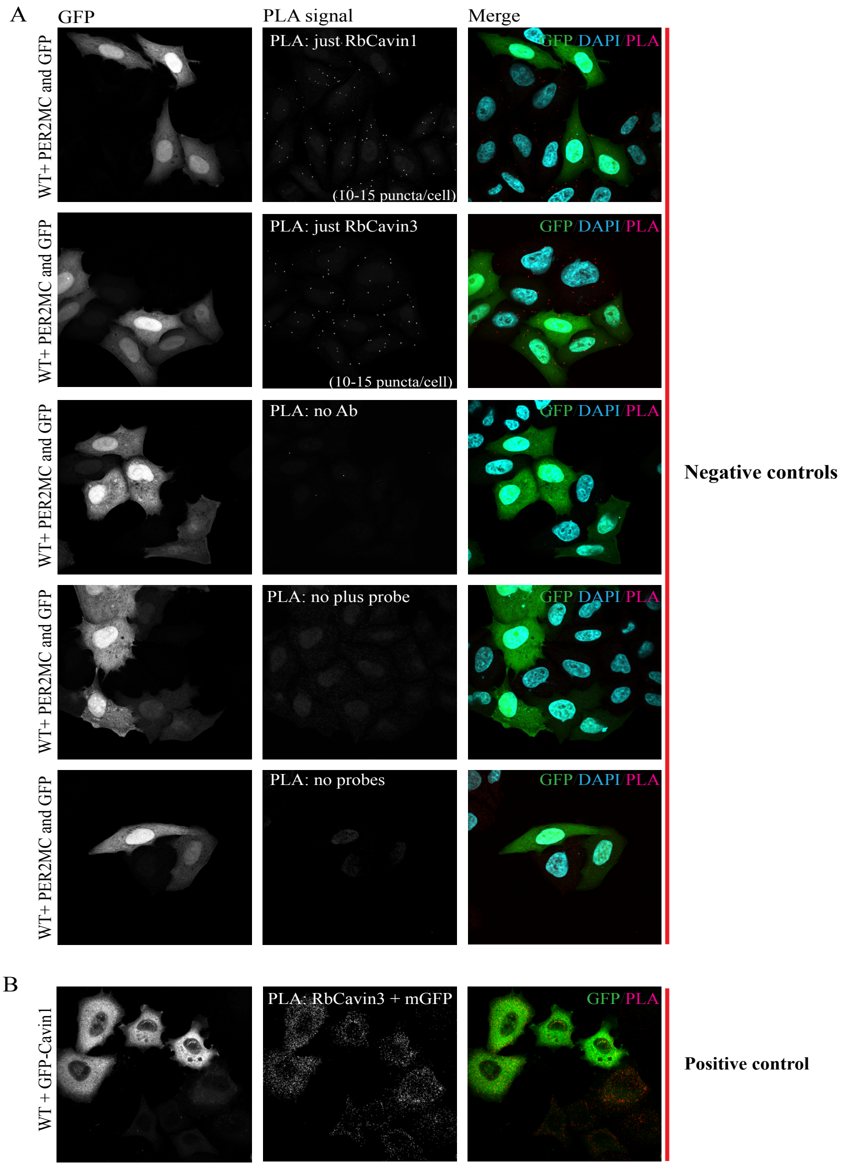


***Supplementary Figure 3*** **/ *PLA controls. A*** */Fluorescence microscopy analysis of PLA signals generated in HeLa cells co-transfected with PER2-MYC and GFP, in the absence of the MYC antibody pair, no primary antibody, missing probe pair and no probes at all.* ***B*** */Positive PLA control using cells transfected with GFP-CAVIN1 and probed with a rabbit CAVIN3 and mouse GFP antibody. Scale bar = 10 µm.*

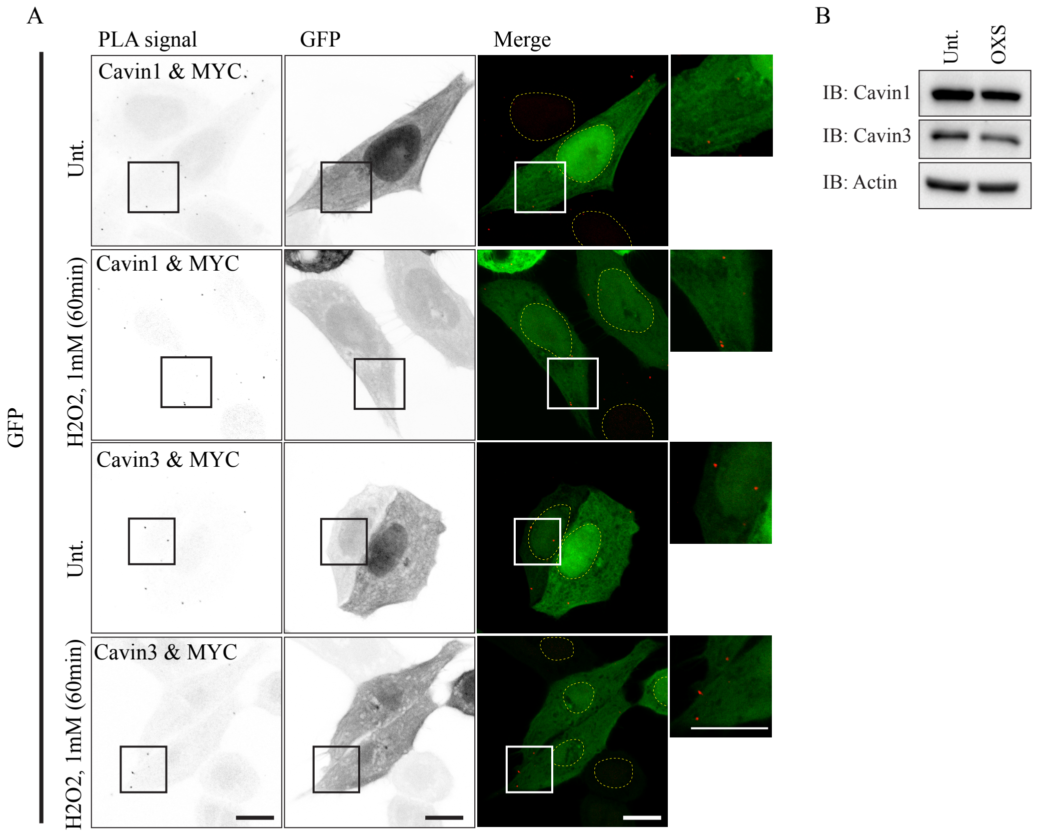


***Supplementary Figure 4** / **PLA controls for oxidative stress. A** /Control cells were transfected with GFP alone (no PER2-MYC) and PLA conducted using a MYC and CAVIN1/3 antibody pair. Low background signal was observed due to absence of PER2-MYC demonstrating assay specificity to the association between PER2-MYC and Cavins. Representative images of PLA signals (inverted image, black puncta), GFP signal and merged images (PLA as red puncta). Dashed circles in indicate the outline of the nucleus. Enlarged images showing the distinct puncta labelling in boxed areas. Scale bar = 10µm. **B** / Immunoblot of endogenous CAVIN1 and CAVIN3 in untreated or oxidative stress (OXS) conditions (1 mM $H_2O_2$, 60min), representative of three independent experiments.*

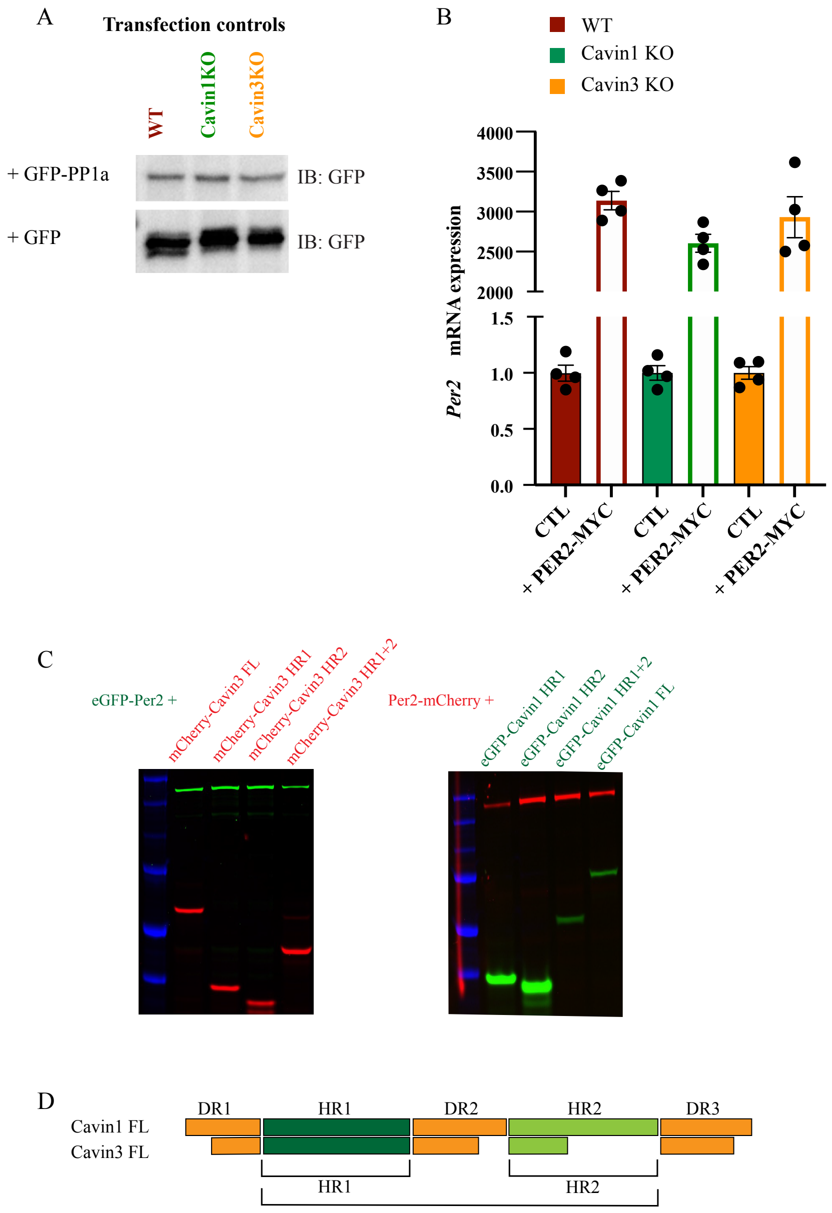


***Supplementary Figure 5** /**CAVIN1 and CAVIN3 KO HeLa cells transfection and expression controls. A** / Immunoblot of transfected PP1α and GFP in WT control and KO cells detected via a GFP antibody, demonstrating similar expression across cell lines. **B** /Cells transfected with PER2-MYC or a GFP control were measured of PER2 mRNA via qPCR to demonstrate similar transcripts across cell lines. Data is presented as fold change to GFP control, mean ± SEM from four independent experiments. **C** /Schematic representation of the helical regions of CAVIN1 and CAVIN3. **D** /Representative in gel fluorescence SDS-PAGE analysis of LTE co-expression of eGFP or mCherry tagged PER2 with domain truncated CAVIN1 or CAVIN3 proteins. In-gel fluorescence shows expression of both components in one lysate. Blot representative of n=3.*

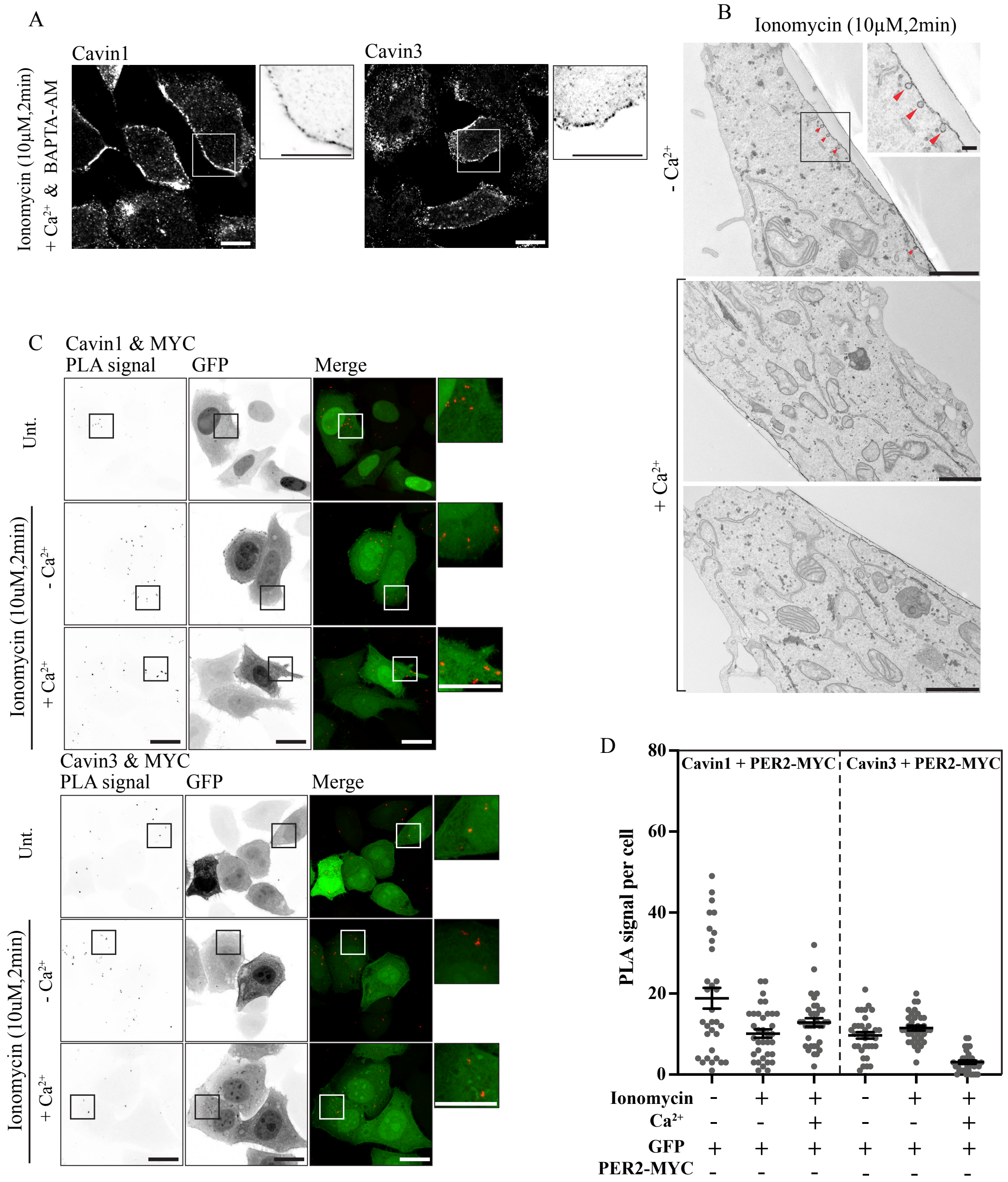


***Supplementary Figure 6*** */* ***BAPTA-AM inhibits ionomycin driven cavin release and PLA controls. A*** */ Representative immunofluorescence images of endogenous CAVIN1 and CAVIN3 labelling in cells with ionomycin with calcium and BAPTA-AM (10 µM ionomycin in 1mM calcium, BAPTA 10 µM, 2min). Inverted enlarged images showing the distinct punctate and diffused labelling.* ***B*** */ Representative EM images of cells treated with ionomycin in the presence (top panel) or absence of* $Ca^{2+}$ *(bottom two panels), the inset is a magnification of the boxed area on top panel. More than 20 images from 6 different areas of two independent experiments were imaged and quantified to assess the caveolae number on the apical surface (caveolae indicated by red arrowheads). Scale bar = 1 µm (0.1 µm inset).* ***C*** */ Control cells transfected with GFP in situ PLA detection of the interaction between PER2-MYC and endogenous CAVIN1 and CAVIN3 in untreated control or ionomycin treatment conditions. Representative images of PLA signals (inverted image, black puncta), GFP signal and merged images (PLA as red puncta). Enlarged images showing the distinct PLA puncta labelling. Scale bar = 10 µm.* ***D*** */ Quantification of PLA signals for the CAVIN1/PER2-MYC and CAVIN3/PER2-MYC interactions, a total of 30-40 cells from three independent experiments. Cells transfected with GFP reporter alone were used as a negative control.*
